# Safety of transcranial focused ultrasound for human neuromodulation

**DOI:** 10.1101/314856

**Authors:** Wynn Legon, Priya Bansal, Leo Ai, Jerel K. Mueller, Gregg Meekins, Bernadette Gillick

**Affiliations:** Division of Physical Therapy and Rehabilitation Science, Department of Rehabilitation Medicine, Medical School, University of Minnesota Minneapolis, MN, USA; Department of Neurological Surgery, School of Medicine, University of Virginia, VA USA; Department of Neurology, School of Medicine, University of Minnesota, MN USA

**Keywords:** Ultrasound, neuromodulation, transcranial, safety, humans, non-invasive brain stimulation, adverse events

## Abstract

**Background:** Low intensity transcranial focused ultrasound (tFUS) is a new method of non-invasive neuromodulation that uses acoustic energy to affect neuronal excitability. tFUS offers high spatial resolution and adjustable focal lengths for precise neuromodulation of discrete regions in the human brain. Before the full potential of low intensity ultrasound for research and clinical application can be investigated, data on the safety of this technique is indicated.

**Objective/Hypothesis:** To provide an initial evaluation of the safety of tFUS for human neuromodulation through participant report and neurological assessment surrounding pilot investigation of tFUS for neuromodulation.

**Methods:** Participants (N = 120) that were enrolled in one of seven human ultrasound neuromodulation studies at the University of Minnesota (2015 – 2017) were queried to complete a follow-up Participant Report of Symptoms questionnaire assessing their self-reported experience and tolerance to participation in tFUS research and the perceived relation of symptoms to tFUS.

**Results:** A total of 64/120 participant (53%) responded to follow-up requests to complete the Participant Report of Symptoms questionnaire. During the conduct of the seven studies in this report, none of the participants experienced serious adverse effects. From the post-hoc assessment of safety using the questionnaire, 7/64 reported mild to moderate symptoms, that were perceived as ‘possibly’ or ‘probably’ related to participation in tFUS experiments. These reports included neck pain, problems with attention, muscle twitches and anxiety. The most common unrelated symptoms included sleepiness and neck pain. There were initial transient reports of mild neck pain, scalp tingling and headache that were extinguished upon follow-up. No new symptoms were reported upon follow up out to 1 month.

**Conclusions(s):** To date, in the literature and including this report, no serious adverse events have been reported as a result of low intensity tFUS for human neuromodulation. Here, we report new data on minor transient events. As currently employed with the parameters used in the studies in this report, tFUS looks to be a safe form of transient neuromodulation in humans.

## Introduction

Transcranial focused ultrasound (tFUS) is a new and promising non-surgical low-energy technique that uses mechanical energy to modulate neuronal activity with high spatial resolution and adjustable depth of focus. tFUS has been used safely and effectively for cortical neuromodulation in mouse [1–4], rat [5,6], rabbit [7], sheep [8], pig [9] and monkey [10,11] models, and has also been demonstrated to be an effective method of transient cortical and sub-cortical neuromodulation in humans [12,13]. In humans, tFUS has been applied to the temporal cortex [14], primary somatosensory cortex (S1) [12,15], secondary somatosensory cortex (S2) [16], primary motor cortex [17,18], primary visual cortex [19] and thalamus [13,20]. tFUS has been shown to affect the amplitude of evoked potentials [7,12,15], the power, phase and frequency of the electroencephalogram (EEG) [12,21]; the blood oxygen level dependent (BOLD) magnetic resonance imaging signal [7,17], as well as tactile [12,15] and reaction time [18] behaviour. In 2014, we found that tFUS targeted at the primary somatosensory cortex in healthy human subjects attenuated the somatosensory evoked potentials generated in the targeted region. These results were specific to the site of neuromodulation and also resulted in a behavioural advantage on somatosensory discrimination tasks [12].

As currently employed, human neuromodulation with tFUS typically involves coupling a single (or multiple [16]) focused single-elements usually in the ~ 250 to 600 kHz range (for efficient energy transfer through skull [22]) to the scalp to target a desired brain region. Transducers for cortical targeting are generally small (~ 30 mm diameter with a ~ 30 mm focal length); produce a ~ 3-4 millimeter lateral and ~1 – 2 centimeter axial resolution, and can be placed anywhere on the head similar to other current methods of non-invasive brain stimulation. Ultrasound is also capable of reaching brain targets deep to the cortical surface as the acoustic waves can be focused to any desired depth within certain limits. Transducers for deep brain modulation are typically larger ( ~ 70 mm diameter) [13,20] to achieve this deeper focal length at reasonable axial resolutions [13]. In addition to adjusting focal lengths, there are a number of parameters that can manipulated when using ultrasound including the acoustic frequency, amplitude, duration, duty cycle, pulse repetition frequency etc. and the efficacy of some of these for successful neuromodulation has been addressed in small animal studies [1,3] though the precise mechanism of acoustic energy for neuronal modulation is largely theoretical [23–25] and the impact of parameter space in humans is not yet well-described. The bioeffects of ultrasound for neuromodulation in humans as described here are likely largely mechanical as opposed to thermal as the parameters used are of low intensity and duration and do not generate sufficient temperatures for thermal modulation [26]. Ultrasound for transient neuromodulation is different from the use of ultrasound for surgery where very high intensities are used to thermally ablate tissues [27,28] or for blood-brain barrier (BBB) opening where high intensities are also used in combination with contrast agents (microbubbles) to intentionally produce cavitation as a means of opening the BBB [29,30].

In its current state, ultrasound for neuromodulation generally follows the safety guidelines of the Food and Drug Administration (FDA) for obstetric diagnostic ultrasound and adult cephalic applications [31]. These include derated limits of spatial peak pulse average (I_sppa_) of 190 W/cm^2^, a spatial peak temporal average of 720 mW/cm^2^ (94 mW/cm^2^ for adult cephalic) and a mechanical index (MI = peak negative pressure/√fc) of 1.9 (MI is an indication of the ability to produce cavitation related bio-effects and can be used as an indication for potential micromechanical damage). These levels of energy are generally respected in human ultrasonic neuromodulation studies even though there are no definitive guidelines for energy deposition into the human brain. There is a long history of ultrasound for diagnostic and therapeutic applications, but explicit expository and dosimetry are still largely lacking [32,33]. There are several thorough reports examining the effect different intensities of ultrasound to affect tissue [34,35] and efforts made to develop thresholds for potential hazards [36,37] though these studies typically use continuous wave schemes and high intensities well beyond the levels used for transient neuromodulation and therefore are not wholly informative for low intensity applications. As such, it is important to assess the safety of ultrasound for human neuromodulation. It is the purpose of this paper to provide an initial assessment of the safety of single element focused ultrasound for human neuromodulation as there is yet any research on participant perceived tolerance and report of symptoms. Here, we report on the findings of a variant of the Participant Report of Symptoms questionnaire [38,39] assessing participants’ perceived tolerance to participation in tFUS and their perceived relation of any symptoms to the ultrasound intervention. Of a group of 120 queried, a total of (N = 64) consented to completing the questionnaire at various time points from immediately post-experiment out to 22 months.

## Material and methods

### Participants

All experiments were conducted with the approval of the Institutional Review Board at the University of Minnesota. A total of 120 volunteer study participants (48 male, 72 female aged 18 – 38 with a mean age of 22.96 ± 2.14 years) provided written informed consent to participate in one or more of the seven experiments from which the data for this study is taken between 2015 and 2017 at the University of Minnesota. Prior to formal experimental procedures, participants were screened via questionnaire for contraindications to non-invasive neuromodulation and none of the participants reported any neurological impairment or identified any contraindications to non-invasive neuromodulation as outlined by Rossi et al. (2009) as identified for transcranial magnetic stimulation [40].

### Experiments

The data for this study is a summary of 64 individual participants that participated in one of seven current or completed experiments conducted in our lab in the Department of Rehabilitation Medicine at the University of Minnesota. Details of the specific objectives or hypotheses of each study are not elaborated upon though details on the tFUS application including transducer specifics, target (cortical, sub-cortical) and parameters (amplitude, duration, etc.) are enumerated. For the purposes of this report, experiments will be referred to by number (1-7) based upon chronological date of commencement. All experiments were conducted in neurologically healthy volunteer participants to test the effect of tFUS on either cortical or sub-cortical [18] neuronal excitability and/or effect on specific behaviours. The environment of the experiments differed as one experiment (Experiment 4) [41] was conducted in a 7T MRI scanner at the Center for Magnetic Resonance Research at the University of Minnesota (https://www.cmrr.umn.edu) and experiment 6 and 7 also involved transcranial magnetic stimulation (TMS) either concurrent with tFUS [18] or as a pre/post measure of motor cortical excitability. For all experiments (except fMRI experiment), participants were seated in a dentist-type chair and asked to either perform a simple task or sit passively for the duration of the experimental protocol. Tasks included a sensory discrimination task [12] and simple stimulus response tasks on a computer. All experiments were repeated measures design with either one of or both an active or passive sham along with the tFUS condition. See Table 1 for experiment recruitment totals and participant demographics.

### Questionnaire and follow-up

For all experiments, participants were retrospectively contacted via email at random time intervals (1 week – 22 months) post experiment for their willingness to participate in a follow-up questionnaire on their experience of undergoing tFUS neuromodulation and perceived relation of any reported symptoms to tFUS. For all experiments participants were contacted via email only once. For experiment 7, participants filled out the questionnaire immediately (~ 20 minutes after tFUS): (This is designated as 0 months in Figures and following text) after experimentation with follow-up at one of four time points post-experiments (1 week, 2 weeks, 3 weeks and 1 month). Those participants who responded affirmatively via email were subsequently contacted via telephone and asked to respond to 20 questions regarding their subjective assessment of their current neurological health (see Supplementary Material for questionnaire). This questionnaire is a variant of the Participant Report of Symptoms questionnaire that has previously been used in other non-invasive neuromodulation studies [38,39]. If there was a positive response to a question indicating perceived experience of the symptom, participants were then asked to rank the symptom severity from 2 - 4 (1 = absent) where 2 = mild, 3 = moderate and 4 = severe. In addition, participants were asked for their subjective assessment of the relation of the symptom to their involvement in the ultrasound experiments. Potential responses were: 1 = unrelated, 2 = unlikely, 3 = possible, 4 = probable and 5 = definite. In instances of positive subjective report – each case was referred to a neurologist (G.M) for medical record review and assessment of reported symptoms.. Participants were remunerated for their participation in this telephone interview session. Total phone call discussion time ranged from 5 – 10 minutes. All phone calls were conducted by one of two lab investigators.

### Transcranial focused ultrasound

For all experiments, the tFUS condition involved acoustically coupling the active face of the ultrasound transducer to the scalp at the pre-determined site depending upon the target of interest. The passive sham condition involved either placing a high acoustic impedance disk on the face of the transducer (Experiments 1 and 2), flipping the transducer over (while on) or simply turning it off during collection (experiment 4 MRI). For active sham conditions, ultrasound was delivered to another brain region (Experiment 3) [18]. Shamming maintained contact of the transducer to the head to mimic the audible sensation of a slight buzzing but attenuate any energy into the head. The audible sound was identical for sham and tFUS conditions and no subjects reported any sensory or perceptual differences between sham and tFUS conditions as previously reported [12,13]. The active sham condition, when employed, involved delivering tFUS to another scalp site (vertex) with the same parameters as the experimental site. The order of sham or tFUS conditions was randomized for each subject.

### tFUS waveforms

All experiments used a single element 0.5 MHz transducer. Transcranial ultrasonic neuromodulation waveforms were generated using a two-channel, 2-MHz function generator (BK 4078B Precision Instruments). Channel 1 was used to gate channel 2 that was a 500 kHz sine wave. Channel 1 was a 5Vp-p square wave burst of 1kHz (N = 500) with a pulse width of 360 μs. This resulted in a 0.5 second duration waveform with a duty cycle of 36%. The output of channel 2 was sent through a 100-W linear RF amplifier (E&I 2100L; Electronics & Innovation) before being sent to the custom-designed focused ultrasound transducer. A total of 3 different transducers were used across the 6 experiments. Experiments 1 and 2 used the same transducer; experiments 3,4 and 7 used the same transducer and experiments 5 and 6 used the same transducer. See Figure 1 for the general ultrasound pulsing strategy and see Table 2 and Table 3 for transducer specifications and stimulation parameters for each study.

**Figure 1.**
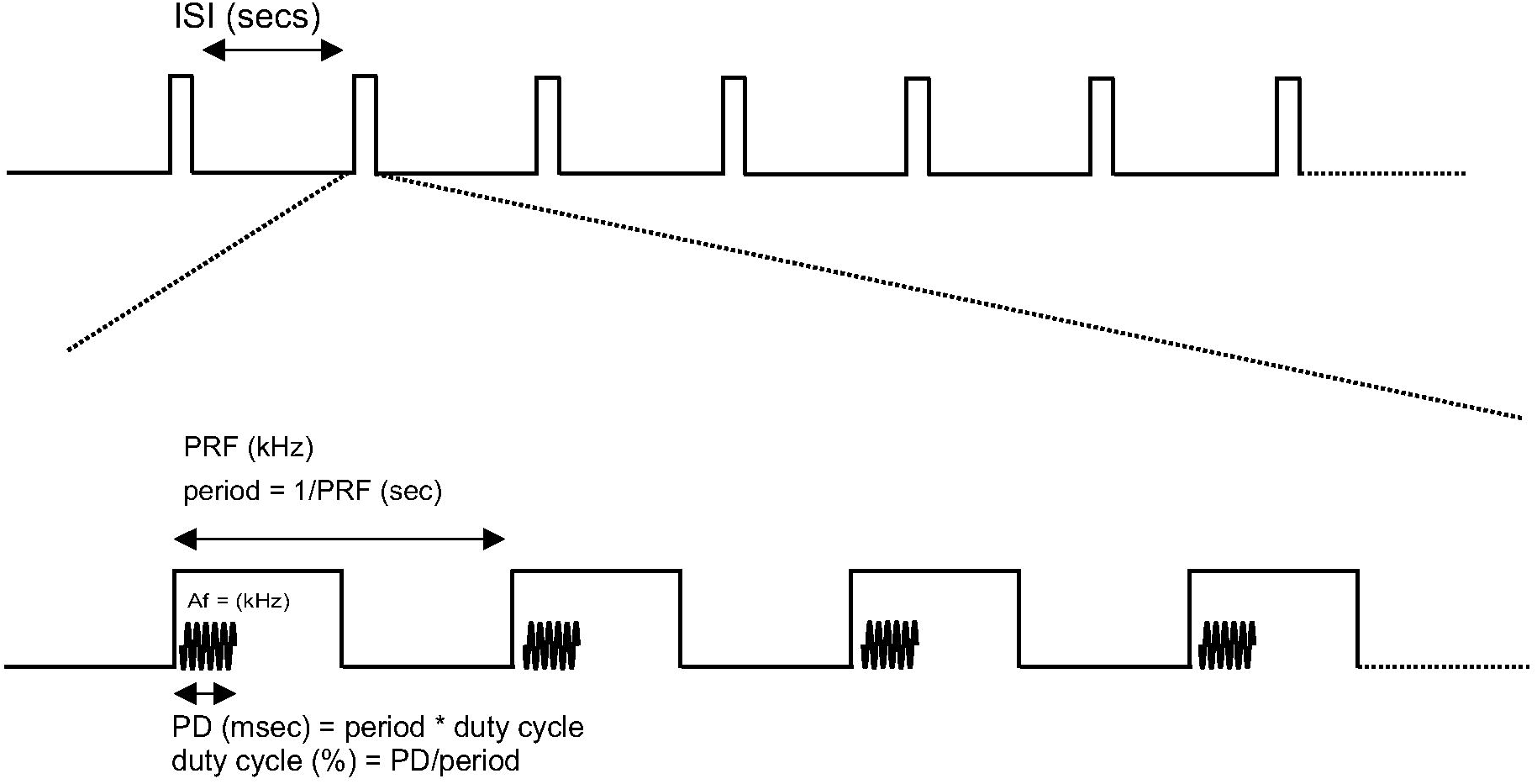
Schematic of ultrasound delivery for human neuromodulation. (Top) Inter-stimulus interval (ISI) in seconds (secs) of delivery of ultrasound. (Bottom) Within each delivery of ultrasound at the given ISI are the parameters that can be adjusted. The frequency of this pulsing is the PRF (pulse repetition frequency in kilohertz (kHz)). Within each ‘pulse’ is the ultrasound or acoustic frequency (Af). The on-time percentage of ultrasound within the period of the pulsing is the duty cycle. This will determine the pulse duration (PD) in seconds and the number of cycles.

## Results

### Response rate

A total of 64/120 (53.3%) participants responded to the email regarding follow-up questionnaire. The mean age of the participants was 22.96 ± 2.14 years (29 Male, 35 Female). See Table 1 for individual experiment demographics and response rates. The time of response post experimental participation ranged from 1 month to 22 months after participation for experiments 1-6 (see Figure 2A). For experiment 7, 17 participants responded to the questionnaire immediately post experiment and then were contacted at a random time post experiment out to one month (Figure 2B).

**Figure 2.**
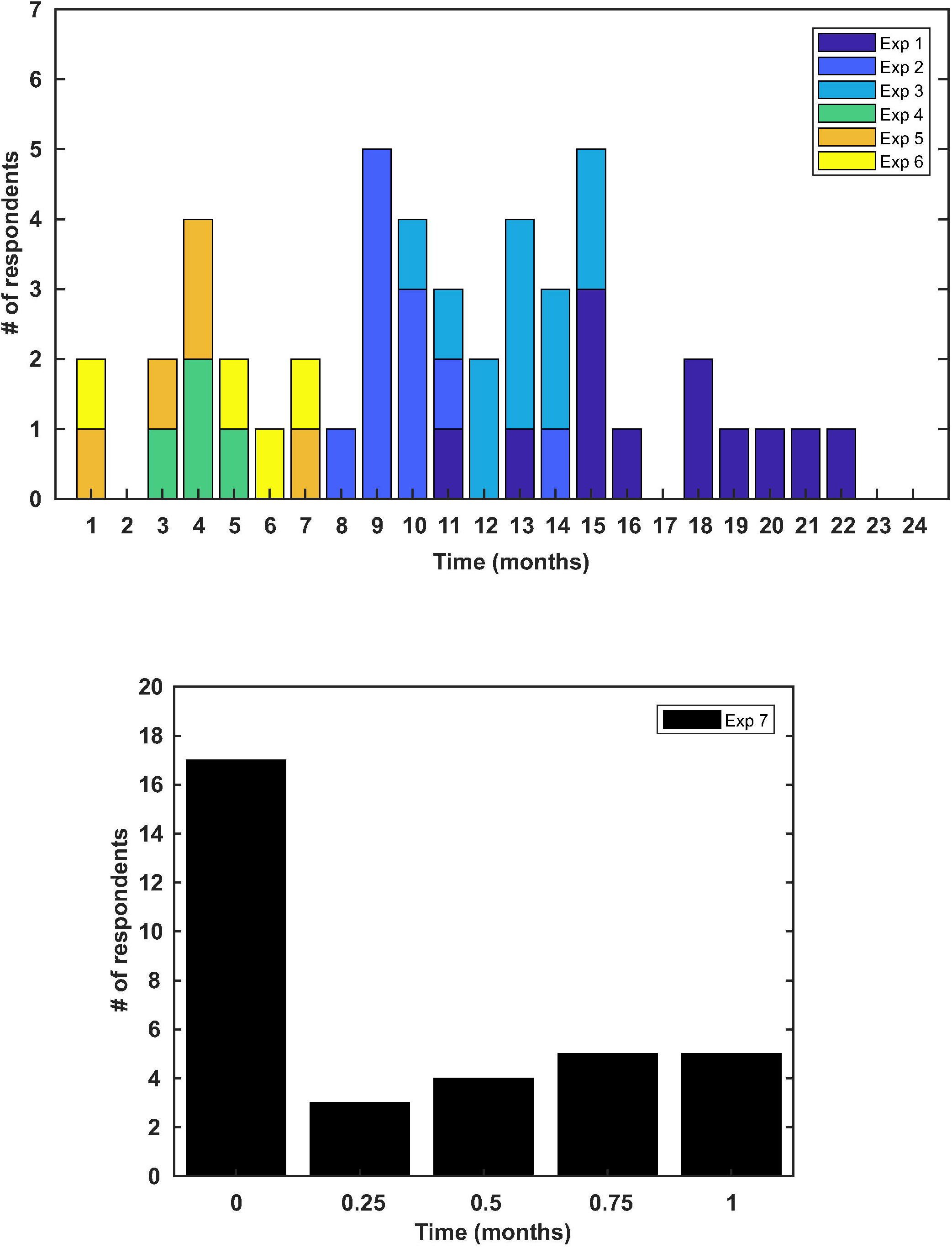
Timeline of respondent follow-up. (Top) Bar graph of the time of questionnaire response of participants from experiments one through six (N = 47) broken down by experiment number. (Bottom) Bar graph of response time for experiment 7. Seventeen participants took the questionnaire the day of the experiment (Time = 0) and responded to the questionnaire again at one of four time points out to one month.

### Symptoms reported

Data from all seven experiments revealed 7/64 reported mild or moderate symptoms that they felt were ‘possible’ or ‘probably’ related to the ultrasound intervention. These included neck pain, difficulty paying attention, muscles twitches and anxiety. There were no reports of any severe or persistent symptoms. Of the other reported conditions, participants rated these as unrelated or unlikely related to the ultrasound intervention (see Figure 3). The most common reported symptom was sleepiness though this was rated as unrelated or unlikely for all instances. Other responses included headache (n = 4), itchiness (n = 5), tooth pain (n = 1) and forgetfulness (n = 4). No participant rated any reported symptom as definitely related to the ultrasound intervention (see Figure 3). There were no qualitative differences in the symptomology between experiments. See Figure 4 for a breakdown of symptoms by experiment for experiments 1-6.

**Figure 3.**
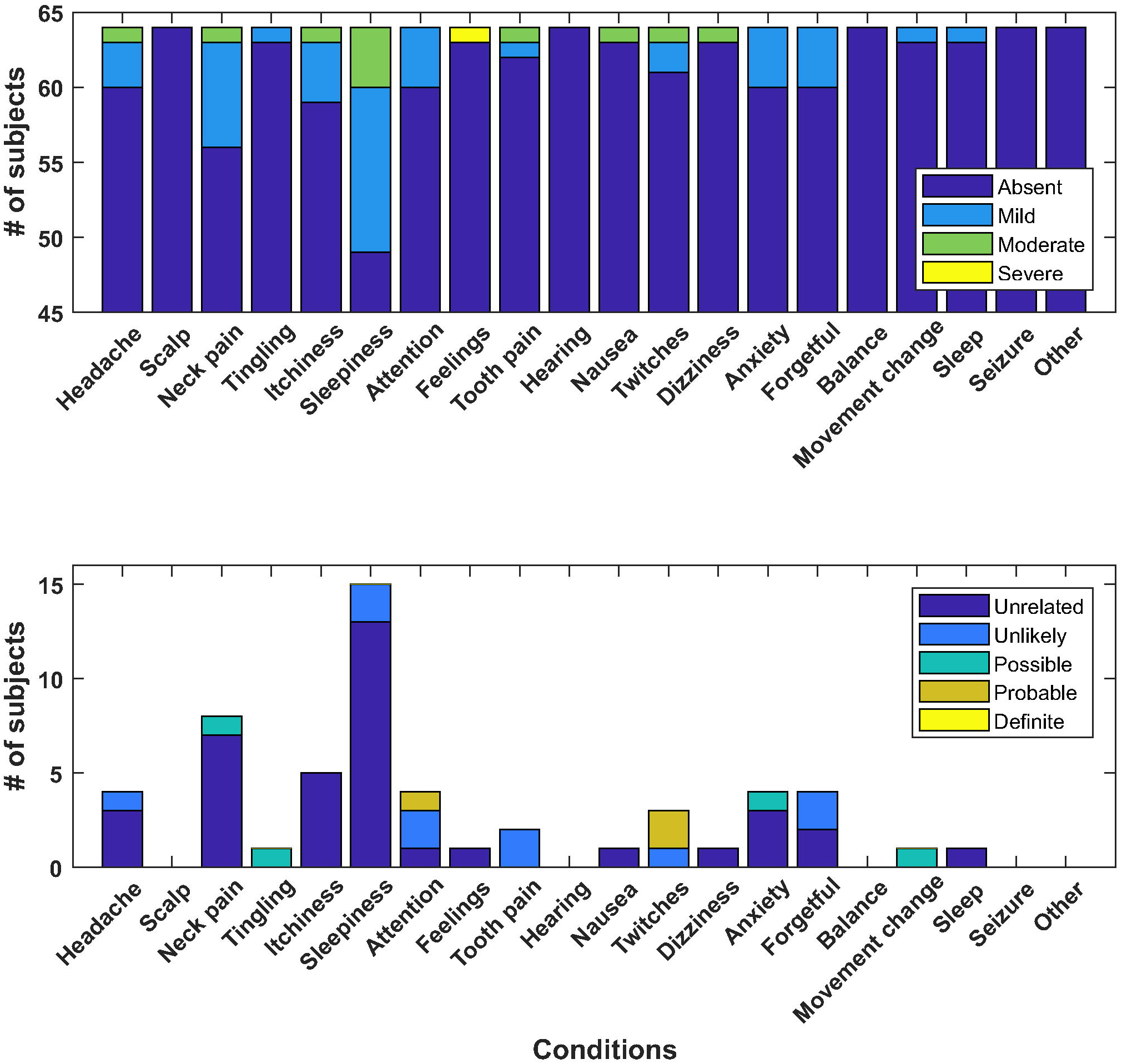
Group report of symptoms. (Top) Total number of responses for all participants (N = 64) collapsed across all experiments (1-7) coded by the severity of the symptom. (Bottom) Total number of responses from all participants (N = 64) collapsed across all experiments coded by the subjective relation of the symptom to the ultrasound neuromodulation intervention.

**Figure 4.**
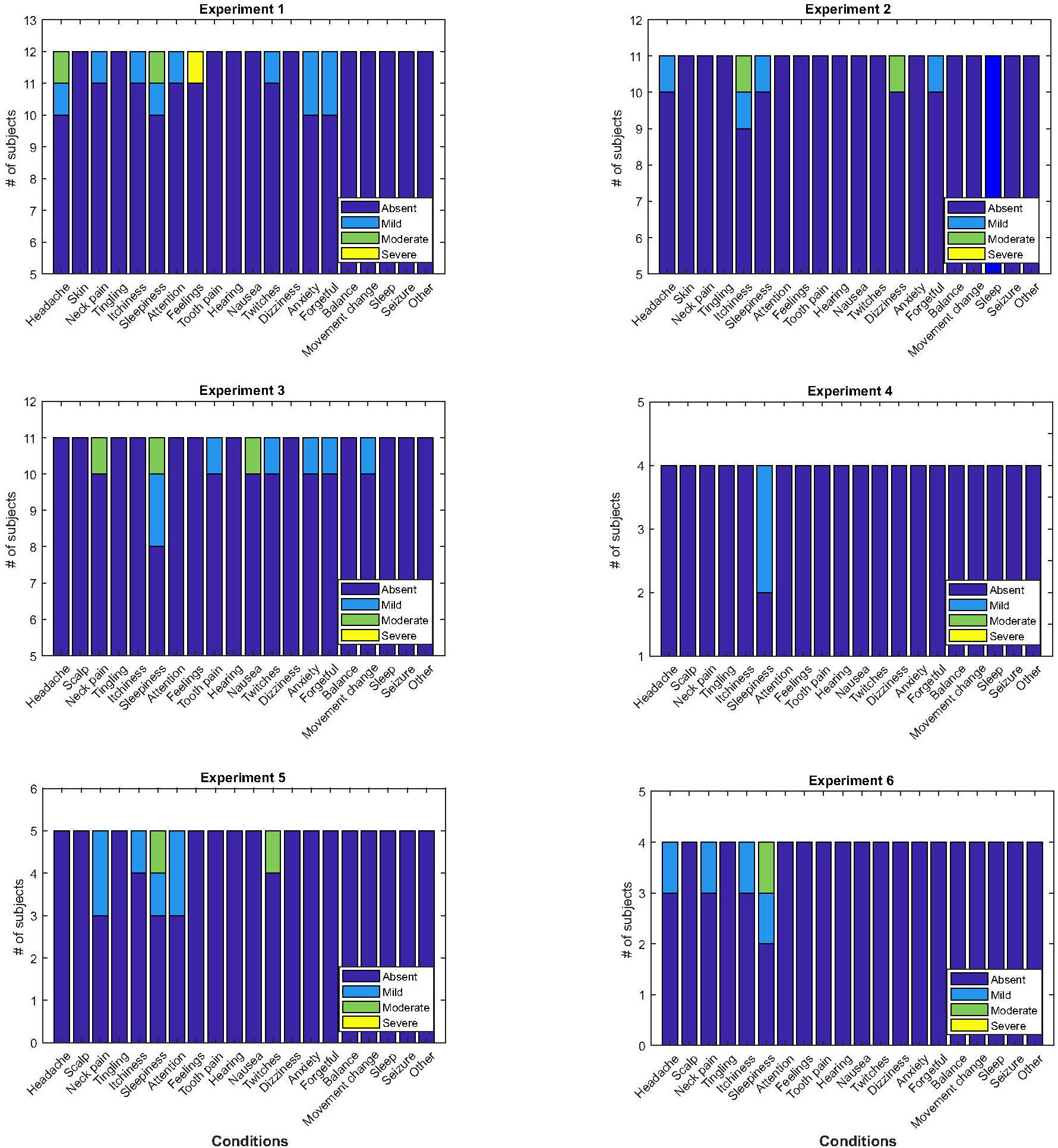
Individual experiment report of symptoms. Individual report of symptoms for experiments 1 – 6.

### Duration of symptoms

In a subset of participants (n = 17) for experiment 7 we collected response to the questionnaire at two time points: immediately following experimentation (~ 20 minutes) and then at a randomly assigned follow-up at 1 week, 2 weeks, 3 weeks or 1 month (see Figure 2B). On day zero, there were three reports of neck pain, three reports of sleepiness, one report of scalp tingling, one report of tooth pain, one report of difficulty paying attention and one report of feeling anxious, worried or nervous and one ‘other’ report of mild back pain (see Figure 5). No participant reported more than one symptom at initial inquiry. Of these reports neck pain was perceived as unrelated in two instances and possible in one. Sleepiness was perceived as unrelated in two instances and unlikely in the other. Tingling of the scalp was perceived as possibly related to the intervention, difficulty paying attention was unlikely, tooth pain was unlikely and anxiousness was perceived as possibly related (Figure 5). At follow-up (1 week to 1 month) these participants did not report any persisting or new effects (Figure 5). Of the 7 participants who did not report any initial symptoms, none reported additional symptoms at follow-up (Figure 5).

**Figure 5.**
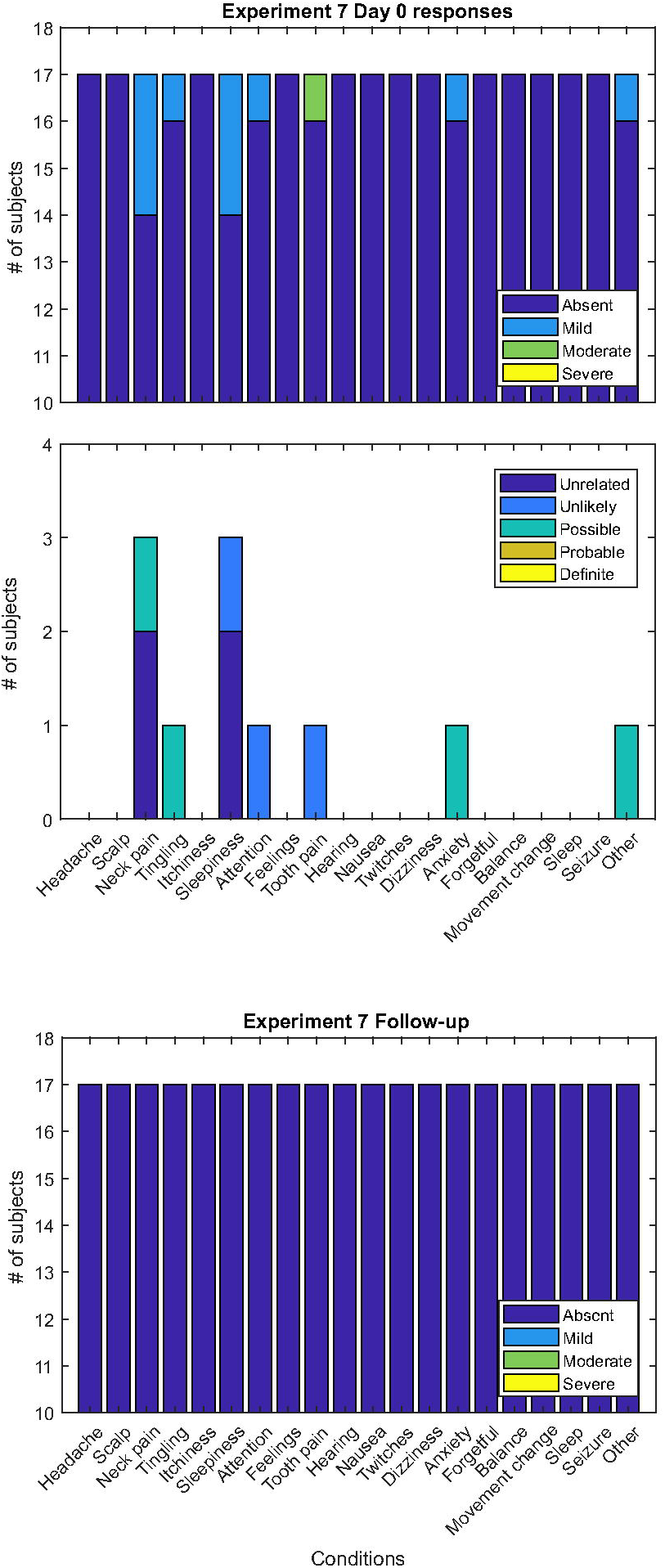
Experiment 7 report of symptoms. (Top) Report of symptoms immediately after completion of ultrasound experiment. (Middle) Perceived relation of immediate symptom to the ultrasound intervention. (Bottom) No new or persistent symptoms were reported at follow-up.

### Correlation of symptom response to tFUS parameters

To gauge the overall positive symptom rate, we tabulated all positive responses regardless of subjective report on the relation to the experimental intervention. The overall positive report of symptoms for all experiments (1-7) included in our neurological questionnaire was 55/1280 total possible positives for an overall positive response rate of 4.3%. Of the 55 total positive responses 38/55 (69%) were judged by the participants to be unrelated to the tFUS interventions, 10/55 (18%) unlikely, 4/55 (7%) possible, 3/55 (5%) probable and 0/55 definitely related. The positive response rates for experiments 1-7 were: 5.4%, 3.2%, 4.5%, 2.5%, 8%, 6.3% and 2.9% respectively (see Figure 6A). The linear correlation of the response rate percentage and mechanical index (MI) was not significant (r = 0.633, p = 0.13) whereas intensity (I_sppa_) was found to have a significant positive correlation with response rate; r = 0.797, p = 0.0319 (Figure 6B).

**Figure 6.**
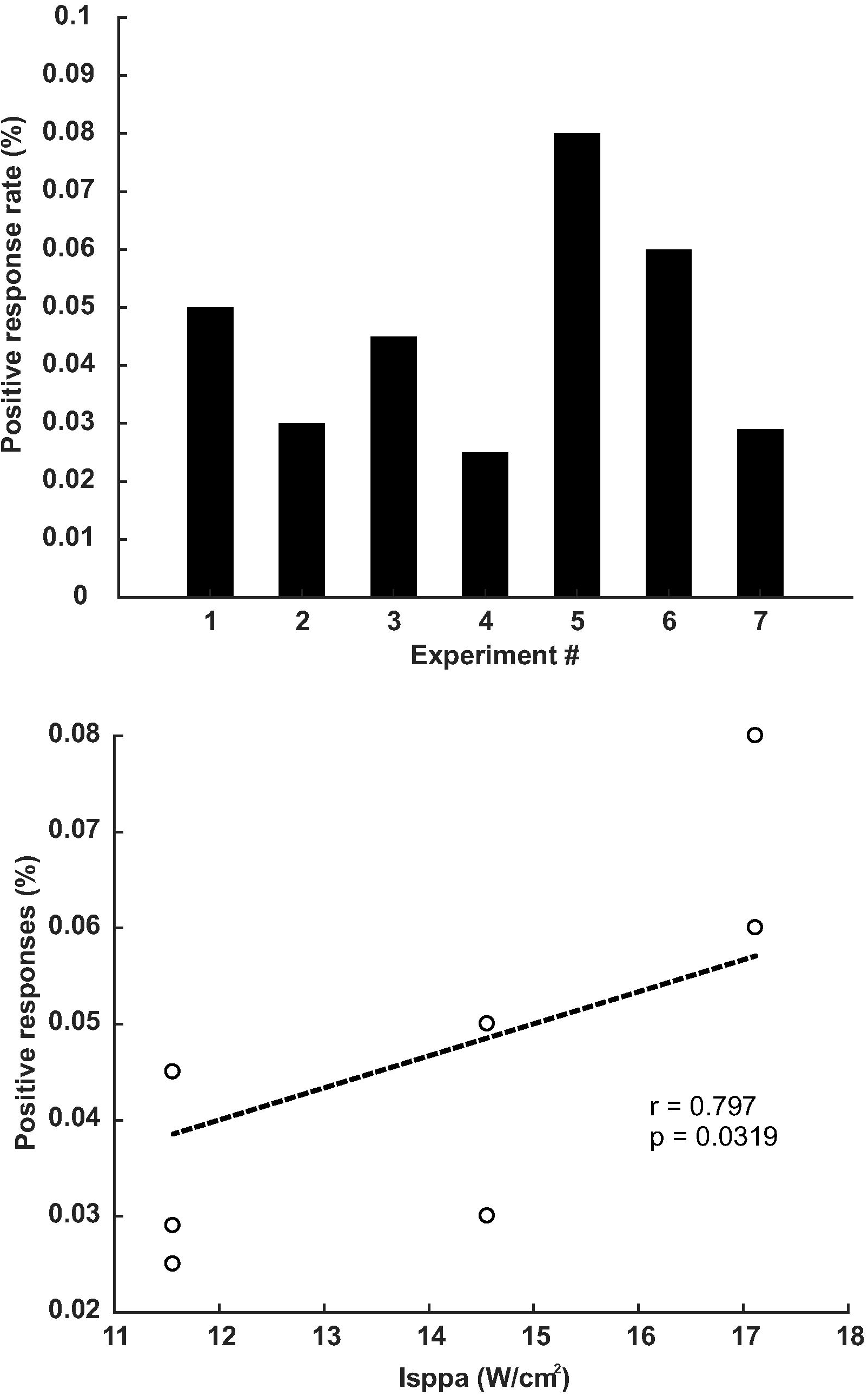
Symptom response rates across experiments. (Top) Symptom response rate for each of the seven experiments regardless of perceived relation to the intervention. (Bottom) Relation of the response rate to the intensity (I_sppa_).

## Discussion

In this report, we provide an initial safety analysis of single element tFUS for human neuromodulation. We collected retrospective data via a participant report of symptoms questionnaire administered over the telephone at varying time points post experiment that ranged from 0 months (day of experiment collected in person) to 22 months post experiment. 64/120 total participants responded to the questionnaire. Symptoms included headache, neck pain, itchiness, sleepiness, problems with attention, tooth pain, muscle twitches, anxiety and forgetfulness. None of these reports were rated as severe and none were reported as definitely related to the tFUS intervention. A subset of participants took the questionnaire immediately after experimentation. Immediate symptoms included mild headache, mild neck pain, and tingling in the scalp. None of these symptoms persisted and no new symptoms were reported upon follow-up out to 1 month. The intensity (I_sppa_) of ultrasound ranged from 11.56 W/cm^2^ to 17.12 W/cm^2^ (in free water) for the experiments included in this study and we found a significant positive linear correlation of the symptom response rate and the ultrasound intensity (I_sppa_). The intensity (I_sppa_) used in these studies is considerably lower than FDA thresholds for ultrasound diagnostics. Despite a lack of definitive causation, and the finding that most of the reported symptoms were believed by the participants to be unrelated to the tFUS intervention, this finding nevertheless speaks to limiting the intensity used in future ultrasound experiments and determining as low as reasonably achievable levels for neuromodulation. Despite the I_sppa_ level being below FDA thresholds, the I_spta_ (spatial peak temporal average) in these studies was above FDA thresholds for diagnostics. I_spta_ is simply the I_sppa_ multiplied by the duty factor providing a metric of the average intensity over the duration of the pulse. One of the main concerns of determining safe intensity levels is estimating the intracranial pressure. The intensities presented here were taken from empirical recordings in free water and hence the derated intensities will be considerably lower. The skull is highly attenuative to ultrasound and the in situ derated pressures are not exactly known but can be estimated using either empirical pressure measurements using a hydrophone through skull fragments or through computer modelling that takes into consideration the acoustic properties of bone and tissue [12,19,42]. For low intensity neuromodulation, in general, ultrasound intensity intracranially is estimated to be attenuated ~3 – 4 fold from values measured in free water [12,13,19] and would thus produce I_spta_ values under the FDA recommended limits for obstetric diagnostics (720 mW/cm^2^) [31].

The experiments documented here had a rather small range of I_sppa_ though used considerably different number of stimulations and different inter-stimulus intervals that would contribute to overall exposure and may influence potential hazard. Indeed, Lee et al. (2016) found that a high number of total stimulations (600) with a low ISI (1 second) resulted in evidence of microhemmorage in sheep despite the intensity being rather low at 6.6 W/cm^2^ [8]. It is currently unclear what constitutes a high number of stimulations or a low ISI though taking these experimental parameters into consideration in the planning of experimental design is prudent. Additional metrics like energy density (J/cm^2^) as has been used in ultrasound sonoporation [43] and neuromodulation parameter [1] studies, as well as the total experimental energy density (J/cm^2^) that takes into account intensity, duty cycle as well as total number of stimulations and the ISI would prove an additional valuable safety metric given the results of Lee et al. (2016) [8] though the relation of total experimental energy density to hazard is not well understood. Experiments examining these assertions are currently being conducted in our lab.

Despite the differences in total number of stimulations and ISI in the experiments reported here, there were no qualitative differences in response rate or in the type of report of symptoms between the seven experiments. The overall symptom response rate and type of symptoms is similar to other forms of non-invasive neuromodulation such as transcranial magnetic stimulation (TMS) and transcranial electric stimulation (TES) [44–48]. In addition to our group, Yoo’s lab has performed human ultrasound neuromodulation studies and completed thorough safety analysis including similar telephone follow-up as well as neurological assessment pre and post experiment including anatomical MRI and reported zero events from their three studies [15,16,19]. In the published literature (N = 233) [12–19,41,49] and including the experiments in this study, a total of 260 individuals have participated in human ultrasound neuromodulation experiments to date with no reported serious adverse events and the data from this report is the first to report on minor transient events associated with the tFUS intervention. Caution is always advised when imparting energy into the brain and further research is recommended examining the effect of total number of stimulations and the inter-stimulus interval and the potential interaction of these parameters with intensity and duty cycle. Consideration of these parameters should be undertaken in experimental design to keep total experimental energy levels as low as reasonably achievable.

## Conclusions

We provide an initial assessment of the safety of ultrasound for human neuromodulation as assessed by participant report of symptom questionnaire. Symptom rate and type are similar to other forms of human non-invasive neuromodulation like TMS and tDCS that have a long standing history of being safe forms of human neuromodulation.

**Table 1 Demographics**

**Table 2 Ultrasound field characteristics**

**Table 3 Ultrasound parameters**

## Acknowledgements

The authors would like to thank Jeff Elias for insightful comments on the manuscript

